# Engineering a Hyper-Adherent *E. coli* Nissle Probiotic Strain that Reduces Intestinal Carriage of *Salmonella* Pathogens in Poultry

**DOI:** 10.64898/2026.06.11.731546

**Authors:** M. Aaron Baxter, Kelsey Greenwood, Kristi L. Anderson, Steven A. Carlson, Bradley D. Jones

## Abstract

Salmonellosis continues to be one of the most important causes of food-borne illness in the U.S. An additional concern with this bacterial pathogen is that infections with multiple-antibiotic-resistant *Salmonella* strains are becoming untreatable infectious diseases. Poultry meat and eggs are major sources of *Salmonella* food-borne illness, due to carriage of these bacterial pathogens in the intestinal microbiome of chickens. A food safety priority, as stated by the USDA, is a significant reduction in carriage of pathogenic *Salmonella* species in poultry which would significantly improve food safety and reduce cases of human salmonellosis contracted from consumption of contaminated poultry. While this goal has been a priority for many years, basic research and animal management efforts have not achieved significant control of *Salmonella* carriage. This study represents an alternative approach to reduce or eliminate carriage of *Salmonella* in poultry flocks. We characterized a type 1 fimbrial allele of *Salmonella* that confers high levels of adherence to various host cells. We then engineered an *E. coli* Nissle 1917 probiotic strain that expresses this *Salmonella* adherence factor at high levels. The *E. coli* Nissle 1917 is used as the scaffold strain for this work since this *E. coli* strain has received the FDA designation of **<u>G</u>**enerally **<u>R</u>**egarded **<u>A</u>**s **<u>S</u>**afe (GRAS) and has been used for many years as a probiotic to treat human intestinal disorders. Our *E. coli* Nissle strain was engineered to use an *in vivo* selection system for a plasmid carrying the cloned *Salmonella* type 1 fimbrial genes, so that the strain can be used as a probiotic without any antibiotic resistance-encoding genes requiring antibiotic selection for maintenance of the desired phenotype. Our probiotic strain displays high levels of adherence to host cells, in fact higher levels of adherence than a *Salmonella* strain carrying the same type 1 fimbrial genes. We demonstrate that the probiotic strain significantly outcompetes pathogenic *Salmonella* strains for adherence to tissue culture cells and *in vivo* experimental challenges revealed that the probiotic strain mediates a significant exclusion of *Salmonella* from the intestines of broilers, layers, and turkeys.

## Introduction

Pathogenic *Salmonella* species are an important cause of foodborne illness in the U.S. with an estimated 1.35 million human infections and 26,500 hospitalizations each year with an associated cost of $ 4.1 billion annually in productivity (https://www.usda.gov/media/press-releases/2022/10/14/usda-releases-proposed-regulatory-framework-reduce-salmonella). Approximately 30% of human cases of *Salmonella* gastroenteritis are linked to poultry consumption making the reduction of pathogenic *Salmonella* serotypes in poultry an urgent food safety priority. Achievement of this food safety goal will substantially reduce human salmonellosis infections and save the associated costs in health and financial expenses. Efforts to identify and characterize the factors of *Salmonella* that allow this pathogen to cause human disease are extensive. Research has identified and characterized genes necessary for cell invasion (SPI-1 genes and associated effectors) [1-4], survival within host cells such as macrophages (SPI-2 genes and associated effectors) [5-7] and survival within the human host that are encoded on additional pathogenicity islands [6, 8, 9] as well as the virulence plasmid [8]. Many other genes have also been studied that encode LPS, serum resistance that allow the bacteria to survive and persist in a host [4, 8]. While these studies yielded enormous information on the biology and virulence strategy of pathogenic *Salmonella* strains in human disease, the impact on the incidence of human salmonellosis has been minimal [10, 11]. Significantly, the *Salmonella* food safety risk begins in the intestinal tracts of poultry, primarily chickens and turkey, where these enteric organisms are important components of the intestinal microbiome. While several metabolic pathways are important for survival and growth in the intestinal environment, the type 1 fimbriae of *Salmonella* play a key role in the ability of these bacteria to persist in the intestinal microbiome of poultry organisms by mediating adherence to poultry intestinal cells [12-17]. In addition to type 1 fimbriae, many *Salmonella* serotypes encode other fimbrial gene clusters (*i*.*e*., - Bcf, Csg, Pef, Lpf, Tafi) although studies to identify roles for these factors in intestinal adherence have largely failed to show any significant role for these adherence factors [18]. In the absence of functional data, these putative fimbriae have no known functions while type 1 fimbriae are essential for the establishment of *Salmonella* in the intestinal microbiome of poultry [12, 13, 17, 18].

*Salmonella* establishes its niche in the intestinal microbiome of poultry by outcompeting other bacteria via type 1 fimbrial adherence to intestinal epithelial cells to gain access to nutrients and space [12, 18, 19]. Type 1 fimbriae of *Salmonella enterica* mediate binding to mannose-containing glycoconjugates on eukaryotic cell surfaces [20, 21]. The fimbriae are encoded by the *fim* gene cluster and are produced on the bacterial surface using the chaperone-usher system of assembly [22]. FimA represents the major structural subunit that is polymerized to form the fimbrial shaft and the adhesin that mediates binding to eukaryotic cells is FimH [19]. There are different alleles of the adhesin (FimH) of *Salmonella* type 1 fimbriae that possess different binding specificities for host cell receptors with some of these alleles having significantly increased binding to host cells [12, 23]. It is presumed that a FimH allele encodes for a highly adherent type 1 fimbriae that provides a competitive advantage to the *Salmonella* serotype for position and growth in the intestinal microbiome, and that different *Salmonella* serotypes have adapted their fimbrial repertoire to exploit different hosts and binding receptors [12, 17, 23, 24].

A prospective study of *Salmonella* strains in poultry supports the idea that *Salmonella* serotypes compete to establish a presence in the intestinal microbiome [25]. This report focused on surveys of carriage of *Salmonella enterica* serovar Gallinarum and *Salmonella enterica* serovar Enteritidis in poultry in England, Wales, and Germany. *S*. Gallinarum, a host-adapted and host-restricted serovar that causes systemic disease and death in domestic and aquatic fowl, was a serious problem in the 1970s that led to a surveillance program that used “test-and-slaughter” to control *S*. Gallinarum in domestic poultry. Over time, this program led to disappearance of *S*. Gallinarum in poultry flocks and, at the same time, the emergence of the human pathogen *S*. Enteriditis as the dominant *Salmonella* serotype colonizing the domestic poultry intestinal tract, with a corresponding dramatic increase of human salmonellosis cases during this same period. The inverse relationship between these two *Salmonella* serotypes led to the conclusion that *S*. Enteriditis filled the ecological niche in the poultry intestinal tract made available by the eradication of *S*. Gallinarum from domestic poultry [25]. A conclusion of this study was that an effective way to reduce carriage of human pathogenic strains in poultry, such as *S*. Enteriditis, would be to engineer a live-oral avirulent strain of *S*. Gallinarum that retained the ability to exclude *S*. Enteriditis from the poultry intestinal microbiome but that had lost its virulence for birds.

This work seeks to define the capacity of an engineered probiotic *E. coli* Nissle 1917 strain expressing hyper-adherent *Salmonella* type 1 fimbriae to selectively colonize the poultry intestinal tract and competitively exclude pathogenic *Salmonella* strains from the gut microbiome. By characterizing enhanced adherence to epithelial surfaces in both tissue culture and poultry intestinal environments, these studies establish the mechanistic foundation for development of a targeted probiotic strategy to reduce *Salmonella* colonization and transmission in commercial poultry production systems.

## Results

### Construction of a nonvirulent *Salmonella* strain that is hyper-adherent

The first attempt to create a nonpathogenic live-oral probiotic strain that might have the potential to reduce *Salmonella* carriage in poultry was to engineer a non-pathogenic *Salmonella* strain that possessed high levels of T1F binding to host cells. This approach was a modification of the approach suggested by Baumler *et al*. [25] which was to create a live-oral avirulent *S*. Gallinarum strain that maintained its ability to exclude pathogenic *Salmonella* strains. We created a noninvasive (SPI-1 mutation), intracellular-survival-defective (SPI-2 mutation) *Salmonella* strain that carries the highly adhesive type 1 fimbriae allele. This *S. enterica* serovar Typhimurium strain is designated as BJ3716 and carries a deletion of the *hilD* and *hilA* <u>u</u>pstream regulatory <u>s</u>equences (URS), which renders *Salmonella* noninvasive [3, 26, 27] and the strain also carries a deletion of the s*saV* gene which is an essential component of the type three secretion system-2 (TTSS-2) machinery [28-30]. Using our standard tissue culture adherence assay, the adherence of BJ3716 was compared to the adherence of pathogenic *S*. Typhimurium SL1344 and a clinical *S*. Typhimurium multiple-antibiotic-resistant DT104 isolate [31] (Figure 1). The negative adherence control for the experiments was S. Typhimurium BJ3714, which is a *fimA* isogenic mutant of BJ3716. The baseline adherence of the nonadherent negative control strain BJ3714 (T1F^-^) was 1.1×10^6^ CFU ± 1.2×10^5^ CFU which represents about 1.1% of the original inoculum that was added to the tissue culture adherence assay well. This level of adherence in a strain that does not express type 1 fimbriae represents nonspecific adherence to the tissue culture cells and the tissue culture plastic. Pathogenic *S*. Typhimurium strain SL1344 had 1.3×10^7^ CFU ± 2.9×10^6^ CFU adherence per well and pathogenic multidrug-resistant DT104 had 9.2×10^6^ CFU ± 6.2×10^5^ CFU per well. In comparison to the negative control strain, this means that 13% of the SL1344 inoculum and 9.2% of the DT104 inoculum added to the adherence assay at the beginning of the experiment adhered in a T1F-specific manner. In direct comparison to the negative control, the two pathogenic *Salmonella* strains, SL1344 and DT104, adhere 13-fold and 9.2-fold better than the non-adherent T1F^-^ BJ3714 strain, respectively. The hyper-adherent *Salmonella* BJ3716 strain had 1.2×10^8^ CFU ± 6×10^7^ CFU adherence per well which is 120% (or 120-fold increase over nonspecific adherence) of the original inoculum added to each well. A percentage above 100% is possible for these quantitative adherence assays since the number of CFU recovered at the end of the three-hour experiment represents both organisms in the original inoculum plus newly divided organisms that adhere to the tissue culture cells. In addition to the levels above the nonadherent strain, BJ3716 adheres 9.2-fold and 13-fold better than the two pathogenic strains SL1344 and DT104, respectively. While the *Salmonella* strain BJ3716 is both nonpathogenic and has a hyper-adherent T1F phenotype, the possibility that a *Salmonella* strain, no matter its genetic characteristics, would be approved as viable probiotic strain for agricultural use seems unlikely. However, an important insight from this first set of experiments is that disruption of the *hilD* and the URS regulatory sequences and *ssaV* have no significant impact on the T1F hyper adherence that we have observed in some *Salmonella* strains. This preliminary data provides impetus to our goal to create a highly adherent, yet avirulent, bacterial strain that will reduce *Salmonella* carriage in poultry. The experiments described above were repeated more than three times; each experiment was performed on a separate day, and samples were plated in triplicate for statistical significance. P values?

**Figure 1.**
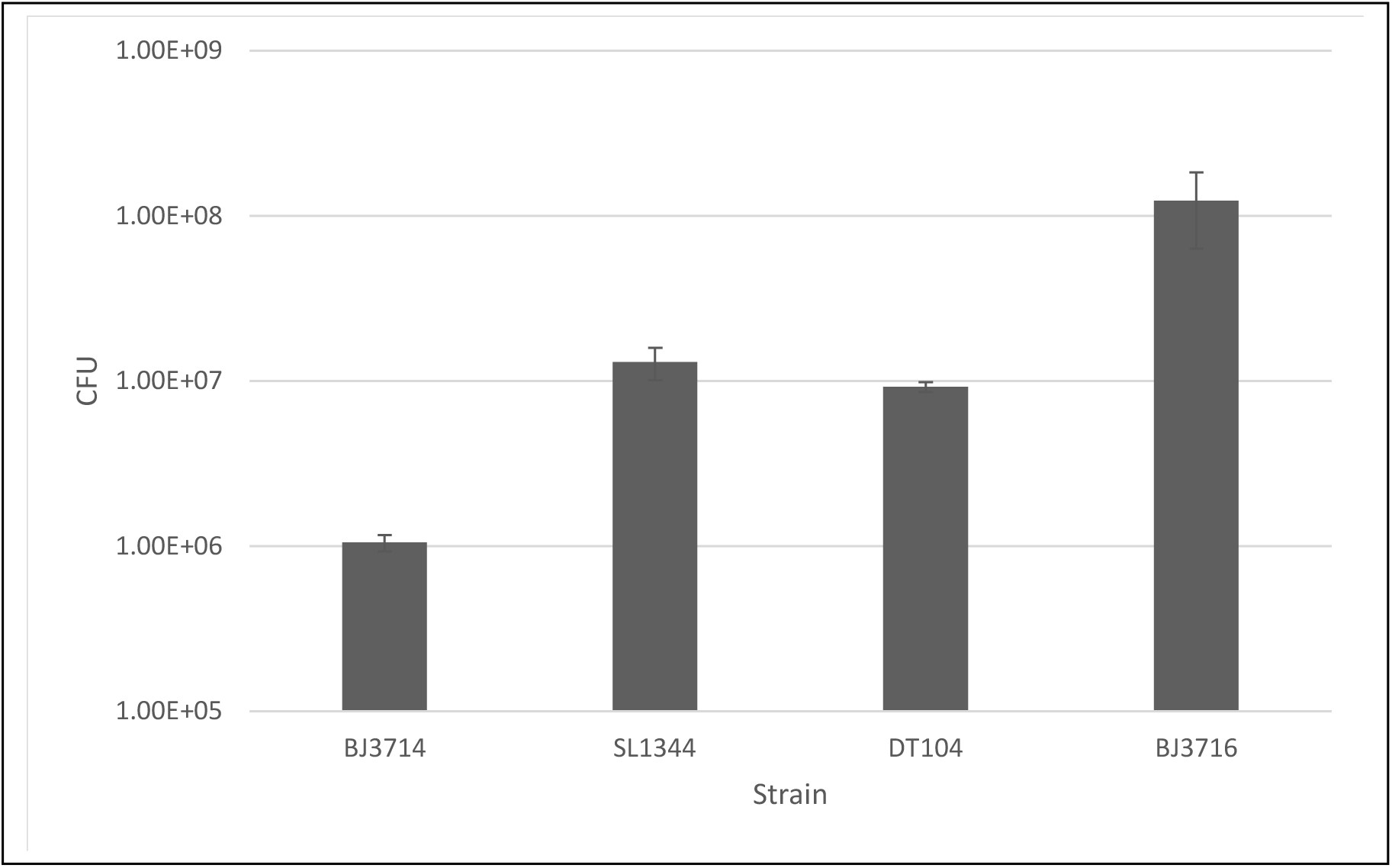
Adherence of *Salmonella* strains to HeLa tissue culture cells *in vitro*. Quantitated adherence is depicted in the bar graph for *S*. Typhimurium BJ3714 (*fimA*-), pathogenic *S*. Typhimurium SL1344 wild type, multiple antibiotic-resistant *S*. Typhimurium DT104, and *S*. Typhimurium BJ3716 Δ*hilD*-URS, Δ*ssaV* hyper adherent allele. Bacteria were grown in conditions that induced maximal expression of adherence by serially passaging cultures that were grown statically for 48 hours. 10^8^ CFU of each strain was added to 10^5^ HeLa cells seeded into the wells of a tissue culture dish and incubated for 3 hours at 37°C in a CO_2_ incubator. Monolayers were washed three times with PBS before lysing the monolayers with 0.2% Triton X-100 in PBS to lyse the tissue culture cells to release attached organisms which were quantitated by diluting samples from each well and plating onto the appropriate agar plate with antibiotics and counting colonies. Experiments were repeated more than three times and statistical significance P value needed.

### Construction of an *E. coli* Nissle 1917 hyper-adherent probiotic strain

Next, we engineered a probiotic in an *E. coli* Nissle 1917 scaffold strain with the highly adherent *Salmonella* T1F genes with the goal of gaining approval for the strain to be approved for use in poultry. Construction of the probiotic strain relies on a pACYC-based plasmid that carries a LT2-derivative type 1 fimbrial gene cluster. This plasmid (pBDJ450) was modified to carry a functional *E. coli* aspartate semi-aldehyde (*asd*) gene which provides *in vivo* selection of the plasmid by complementing an *asd* mutation on the *E. coli* Nissle chromosome. This plasmid, designated pBDJ451, carries a functional *Salmonella* type 1 fimbrial gene cluster and the functional *asd* gene. Since *E. coli* Nissle produces an *E. coli* version of type 1 fimbrial gene, we deleted the *fimH* gene of the *E. coli* T1F to make the *E. coli* Nissle scaffold strain nonadherent. The phenotype of this intermediate strain was confirmed by agglutination test to be nonadherent. Since the final probiotic strain should not carry an antibiotic resistance gene, we took advantage of an *in vivo* selection system based upon the *asd* gene for selection of plasmid pBDJ451. The *E. coli* Nissle 1917 *asd* gene converts aspartate into diaminopimelate (DAP), an essential unique amino acid in the *E. coli* bacterial cell wall that cannot be scavenged from eukaryotic organisms. The chromosomal *asd* gene was deleted using the one-step inactivation procedure [32] which replaced the functional *asd* gene with a chloramphenicol resistance gene, flanked by FRT sequences. The chloramphenicol resistance gene was subsequently deleted using the Flp recombinase on the temperature-sensitive plasmid pCP20, which created an *E. coli* Nissle Δ*asd* T1F^-^ mutant strain. The mutant was unable to grow without the addition of DAP to the growth media but when pBDJ451 was transformed into *E. coli* Nissle 1917 Δ*asd* the strain grew as well as wild type *E. coli* Nissle without the addition of DAP, indicating that the *in vivo* selection proceeded as expected.

### Adherence phenotypes of *E. coli* Nissle 1917 pBDJ451

After construction of the hyper-adherent probiotic strain, we confirmed that *E. coli* Nissle 1917 Δ*asd* pBDJ451 expressed mannose-sensitive *Salmonella* type 1 fimbriae using a standard agglutination assay. *E. coli* Nissle 1917, *E. coli* Nissle Δ*asd* pBDJ451, and *S*. Typhimurium BJ3716 were grown in conditions that induce expression of type 1 fimbriae. Each of the three strains were tested for agglutination in the presence or absence of mannose or in the presence of anti-*Salmonella* type 1 fimbriae antibody, which inhibits *Salmonella* type 1 fimbriae-mediated agglutination [33, 34]. As seen in Figs. 2A and 2B, *E. coli* Nissle 1917 agglutinates yeast cells in a mannose-sensitive manner but the agglutination mediated by the type 1 fimbriae produced by *E. coli* Nissle 1917 is not blocked by the anti-*Salmonella* type 1 fimbriae antibodies (Fig. 2C), indicating that the agglutination observed for *E. coli* Nissle 1917 wild-type is mediated by *E. coli* type 1 fimbriae. Similarly, the hyper-adherent *Salmonella* BJ3716 strain agglutinates yeast cells in a mannose-sensitive manner (Figs. 2G and 2H) but, in contrast to *E. coli* Nissle agglutination, is blocked by the anti-*Salmonella* type 1 fimbriae antibodies (Fig. 2I). The probiotic *E. coli* Nissle Δ*asd* pBDJ451 also displayed a strong yeast agglutination phenotype that was blocked by mannose (Figs. 2D and 2E) but importantly the *Salmonella* anti-type 1 fimbriae antibodies inhibited the agglutination reaction (Fig 2F) confirming that the *E. coli* Nissle Δ*asd* pBDJ451 strain produces significant amounts of *Salmonella* type 1 fimbriae that bind to eukaryotic cells in a mannose-sensitive manner.

**Figure 2.**
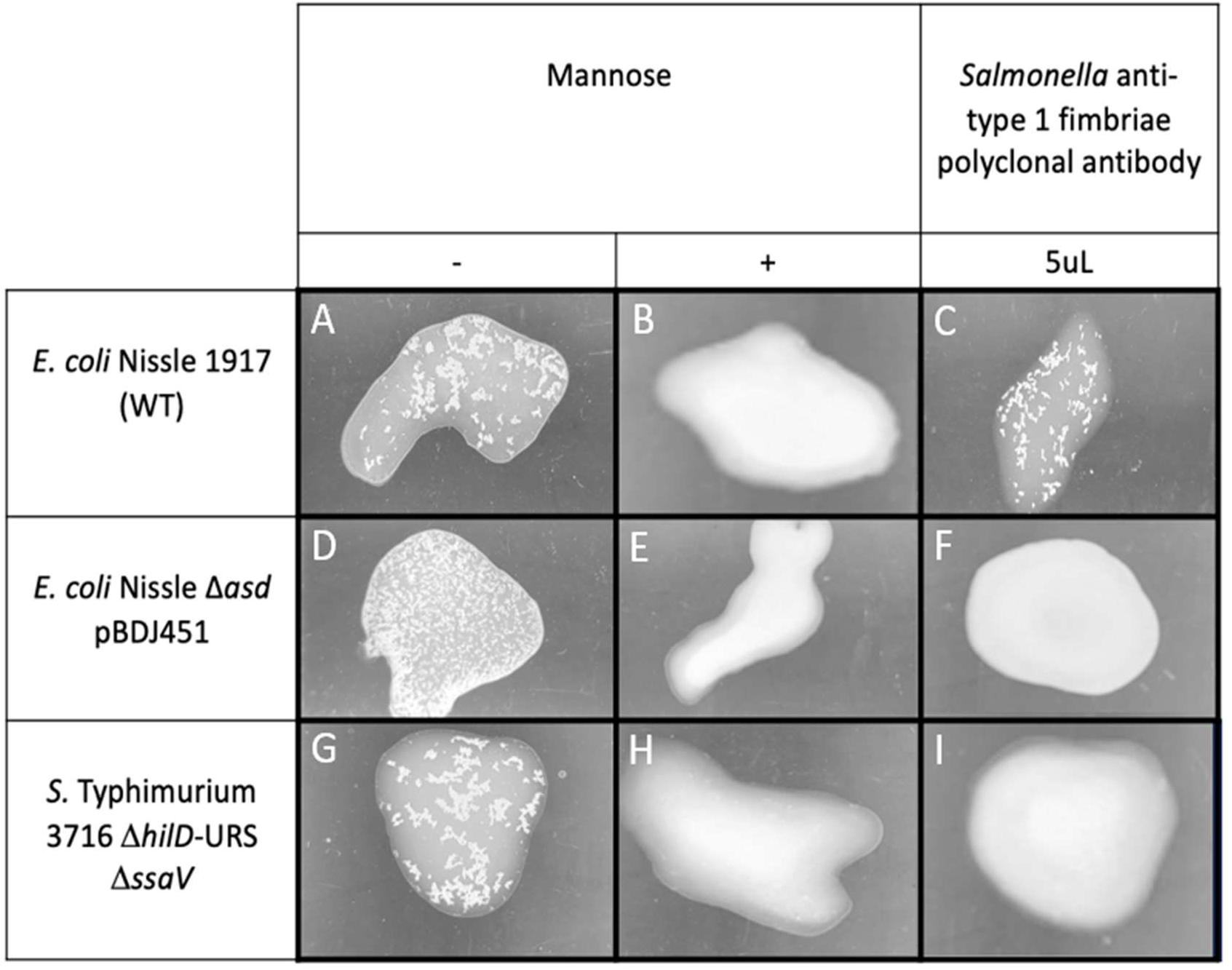
Agglutination phenotype of the *E. coli* Nissle 1917 pBDJ451 probiotic strain. The *E. coli* Nissle pBDJ451 probiotic strain, the *E. coli* Nissle 1917 control and the *S*. Typhimurium BJ3716 control were grown in adherence inducing conditions. Typically, yeast cells were prepared fresh on the day of the assay and 20 μl of yeast cells were mixed with 20 μL of 10-fold concentrated bacterial cultures and deposited on a clean microscope slide. The mixture was gently mixed for 2-5 minutes until a positive control strain agglutinated. As seen in panels A, D, and G, each of the strains clumped dramatically, indicating a positive reaction. In all instances, 1 mM mannose inhibited the agglutination observed in the absence of mannose (panels B, E, and H). *Salmonella* anti-type 1 rabbit polyclonal antibody specifically inhibits *Salmonella* type 1 fimbriae activity but not *E. coli* type 1 fimbriae activity as seen in panels F and I but in panel C.

We previously demonstrated that *S*. Typhimurium strain BJ3716 possessed adherence that was 10-fold higher than either *S*. Typhimurium SL1344 or *S*. Typhimurium DT104 in tissue culture adherence assays (Figure 1). We wanted to quantitate adherence produced by *E. coli* Nissle 1917 Δ*asd* pBDJ451 and compare it to adherence observed with *S*. Typhimurium strain BJ3716. In a comparative tissue culture adherence assay, we observed 1.0×10^8^ CFU ± 4×10^7^ CFU adherence for *S*. Typhimurium BJ3716 and 5.2×10^8^ CFU ± 9×10^7^ CFU for *E. coli* Nissle 1917 Δ*asd* pBDJ451 (Fig. 3). Thus, the *E. coli* Nissle 1917 pBDJ451 probiotic strain displayed even higher adherence (5.2-fold higher) than the hyper-adherent *S*. Typhimurium strain BJ3716. This surprising result is likely due to *Salmonella* type 1 fimbrial proteins being produced by multicopy plasmid pBDJ451 and bodes well for the ability of this strain to outcompete pathogenic *Salmonella* strains in binding to host cells in the intestinal environment. These experiments were repeated three times, each experiment being conducted on a separate day and samples were plated in triplicate for statistical significance. P values?

**Figure 3.**
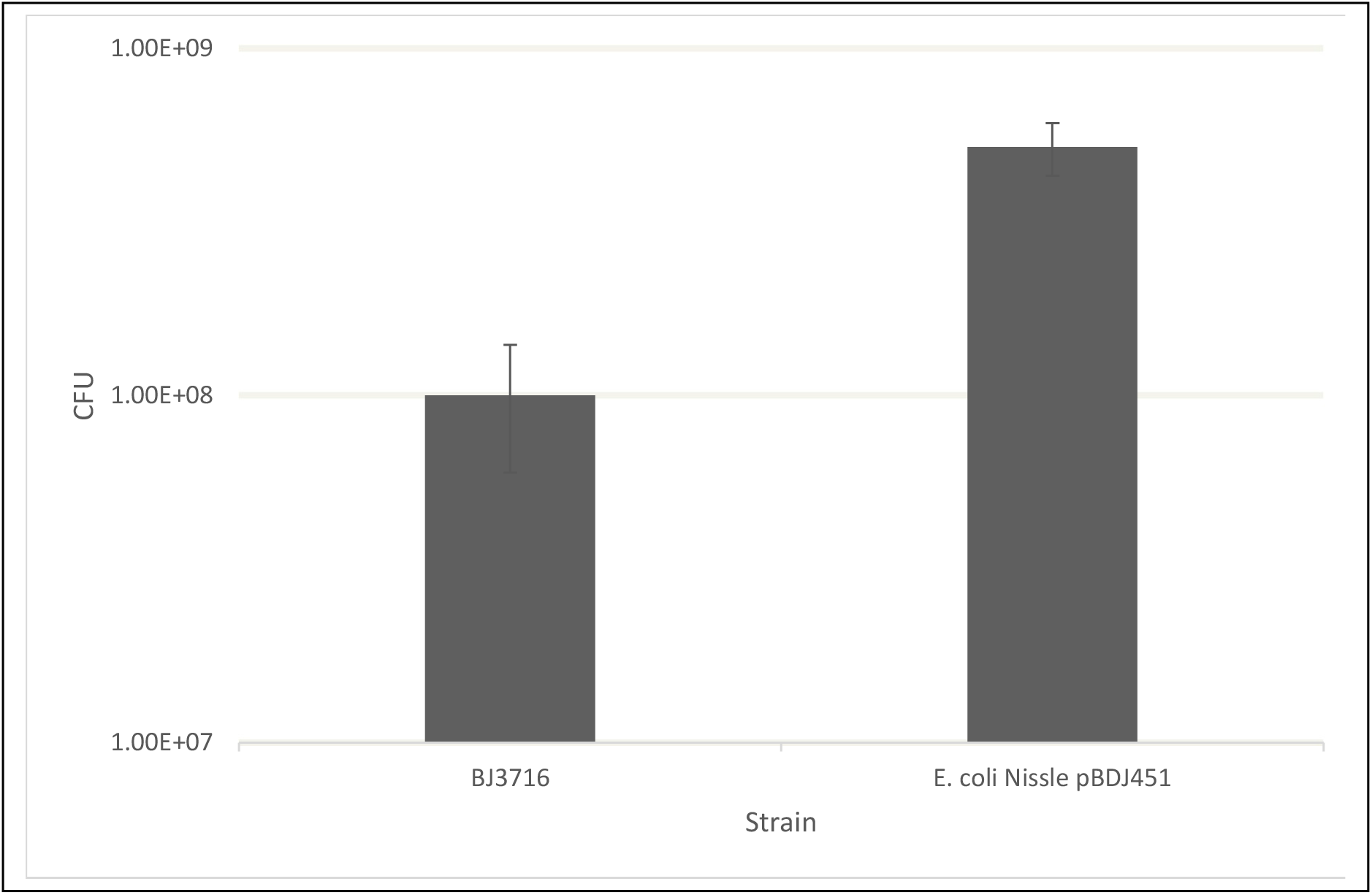
Adherence of hyper adherent *S*. Typhimurium BJ3716 compared to *E. coli* Nissle 1917 pBDJ451. Adherence of the *E. coli* Nissle 1917 pBDJ451 was quantitated using the standard adherence assay and compared to hyper adherence observed for BJ3716. The adherence observed in the *E. coli* probiotic strain was ∼5-fold higher than observed for BJ3716. Experiments were repeated three times, p value needed.

### Competitive adherence assay as a model for competitive exclusion

We demonstrated that the engineered *E. coli* Nissle 1917 Δ*asd* pBDJ451 recombinant strain possesses the ability to bind to tissue culture cells at high levels which is ∼500-fold higher than nonadherent *Salmonella* strain BJ3714 *fimA* mutant, 20-50-fold higher adherence than pathogenic *Salmonella* strains SL1344 and DT104, respectively (Fig. 1), and even ∼5-fold higher than engineered hyper-adherent *Salmonella* strain BJ3716 (Fig. 3). We wanted to assess whether there is a difference in binding activity when two strains were competing to adhere to the same tissue culture monolayer. We developed this assay as a model to assess one aspect (adherence) of the ability of the *E. coli* Nissle 1917 pBDJ451 probiotic strain to competitively exclude pathogenic *Salmonella* strains from binding to host cells. Adherence assays were performed as described for single strain adherence assays with the exception that both strains (1×10^8^ CFU/well) were added to the well at the same time and allowed to bind to the tissue culture monolayer. As shown in Fig. 4 (left-side bars), the hyper-adherent *E. coli* Nissle pBDJ451 probiotic strain adhered 144x higher to tissue culture cells when competing with *Salmonella* SL1344 and 185x higher when competing with *Salmonella* DT104 for adherence (Fig. 4, right-side bars). These results are very encouraging and suggest that the *E. coli* Nissle 1917 pBDJ451 has the potential to adhere at very high levels to chicken intestinal epithelial cells and exclude pathogenic *Salmonella* strains from adhering at levels that are typical for *Salmonella* when colonizing the chicken intestinal tract. The competition assays were repeated a total of three times on three separate days with the data shown in Fig.4 representative of each of the experiments. Samples were plated in triplicate for each individual experiment as well to provide statistical significance. P values?

**Figure 4.**
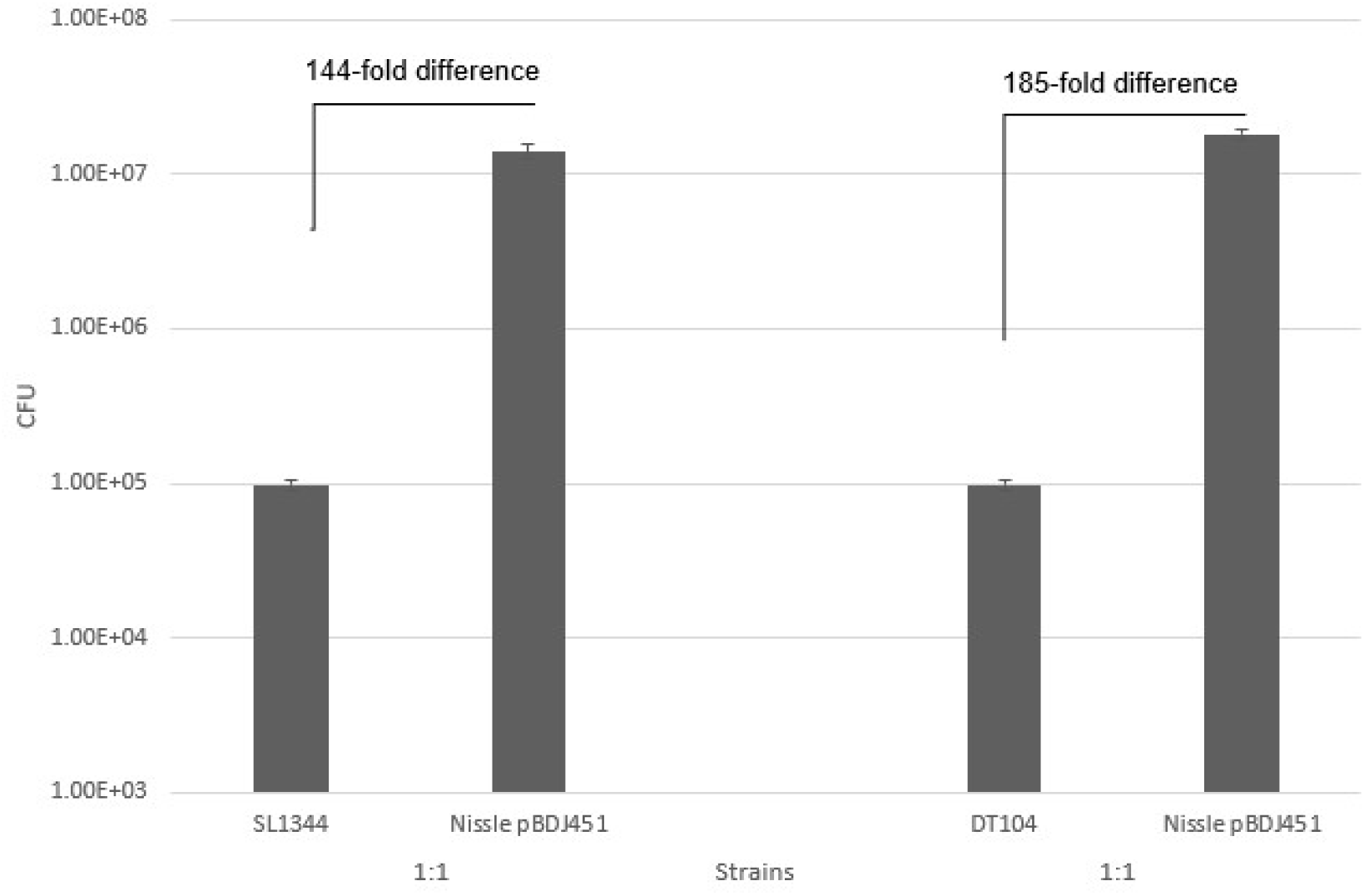
Competitive adherence assay examining the adherence of the *E. coli* Nissle pBDJ451 probiotic strain in the presence of pathogenic *Salmonella* strains. The ability of strains to competitively adhere to tissue culture cells was measured in a standard adherence assay. In one set of experiments, the binding of *E. coli* Nissle pBDJ451 was compared to *S*. Typhimurium SL1344 binding (left-side bars) and the other comparison was *E. coli* Nissle 1917 pBDJ451 compared to *S*. Typhimurium DT104 (right-side bars). Each strain was grown in conditions that induce the adherence phenotype. 10^8^ CFU of each strain was added to tissue culture wells seeded the previous day with 10^5^ HeLa cells and adherence to the cells was allowed to proceed for 3 hours. Adhered bacteria were washed three times with PBS, the cell monolayers were lysed with 0.2% Triton X-100 in PBS, diluted with L broth and then dilutions from each well were plated on agar plates with selective antibiotics for each strain. Experiments were repeated more than three times and the data shown is a best representation of the reproducible results. P value needed.

### *E. coli* Nissle 1917 pBDJ451 competitively excludes *Salmonella* in poultry

To determine if *in vitro* hyper-adherence leads to competitive exclusion of wild-type *Salmonella* from the intestines of poultry, *E. coli* Nissle 1917 pBDJ451 was orally inoculated into birds for three consecutive days. On the fourth and fifth days, birds were orally inoculated with wild-type *Salmonella*. Cloacal swabs were then obtained every 3-4 days and swabs were subjected to enumerative *Salmonella* culture. At the end of the study birds were euthanized and ceca were harvested and subjected to enumerative *Salmonella* culture. As a control, a parallel cohort of birds were orally inoculated with parent *E. coli* Nissle 1917. For studies involving layers, eggshell contamination and oviduct colonization of *Salmonella* were also quantitated. Fold reduction was calculated as (CFU/gm for *E. coli* Nissle 1917 pBDJ451-inoculated birds)/(CFU/gm for *E. coli* Nissle 1917-inoculated birds) whereby 100 indicates a complete elimination of *Salmonella*.

As shown in Fig. 5, cloacal shedding and cecal colonization of *Salmonella* Typhimurium was significantly reduced in broilers, turkeys, and layers inoculated with *E. coli* Nissle 1917 pBDJ451. *Salmonella* egg shell contamination and ascending colonization of the oviduct was also significantly reduced in layers orally inoculated with *E. coli* Nissle 1917 pBDJ451. This reduction in *Salmonella* burden was also observed for layers orally inoculated with *Salmonella* Enteriditis.

**Figure 5.**
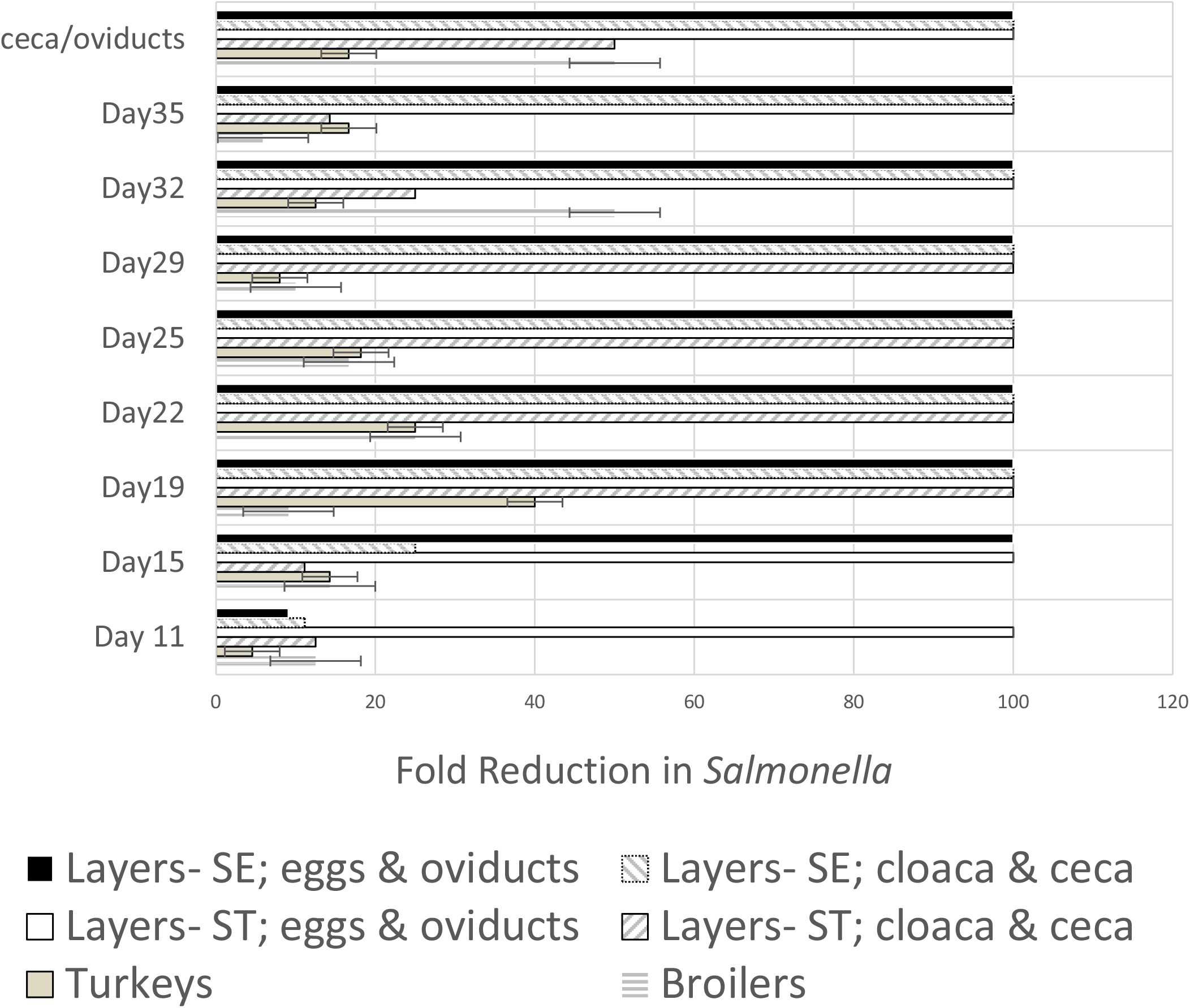
*In vivo* reduction of cloacal shedding, cecal colonization, egg contamination, and oviduct colonization of experimentally introduced *Salmonella* in poultry inoculated with *E. coli* Nissle 1917 pBDJ451. Broilers, layers, and turkeys were orally inoculated with *E. coli* Nissle 1917 pBDJ451 followed by oral inoculation with *Salmonella. Salmonella* burden (cloacal shedding for all birds and egg contamination for layers) was measured every three to four days on eight occasions. At the end of the study, birds were euthanized and ceca were harvested from all birds and oviducts were harvested from layers for enumerative *Salmonella* culture. All three types of birds were inoculated with *S*. Typhimurium (ST) while one layer experiment involved inoculation with *S*. Enteriditis (SE). As a control, birds were orally inoculated with *E. coli* Nissle 1917 and fold reduction of *Salmonella* was calculated based on the presence of *Salmonella* in the control birds whereby a value of 100 indicates complete elimination of the *Salmonella*. Data represent the mean ± sem for 20 birds per treatment group.

## Discussion

The Centers for Disease Control and Prevention estimate that *Salmonella* species cause an estimated 1.35 million cases and 26,500 hospitalizations per year in the United States [35]. More than 30% of these cases are the result of consumption of poultry that is contaminated with pathogenic *Salmonella* species. Chicken and turkey carcasses are commonly contaminated with *Salmonella* species, as well as other intestinal organisms, and fecal contamination of eggs can also be a major source of microbiologic contamination. *S*. Typhimurium and *S*. Enteritidis are commonly isolated serovars isolated from contaminated chicken and often cause symptomatic clinical disease. While the food safety problems associated with *Salmonella* carriage in poultry are well understood because of extensive research efforts, providing solutions to the issue have remained an ongoing effort. A variety of approaches are being used to reduce the risk of *Salmonella* carriage in poultry that include the use of prebiotics, probiotics, bacterial subproducts, bacteriophage therapy and vaccines that provide *Salmonella* control [36]. Some of these approaches are promising, but each has various difficulties or shortcomings that need to be addressed.

The probiotic approach described in this study seeks to take advantage of *Salmonella* adherence factors that are crucial to the ability of the bacteria to colonize the poultry intestinal tract. Research efforts, by many groups, seeking to identify and characterize the key factors that allow *Salmonella* to establish colonization in poultry have led to the general conclusion that type 1 fimbriae are a key factor in colonizing the poultry intestinal tract [12, 13, 17, 18]. A variety of studies have characterized the genetics, expression, and functions of these organelles [14, 19, 22, 37-40]. Other work has explored the host immune response to *Salmonella* fimbriae. Efforts have been focused on determining if host immunity produces a neutralizing response to fimbrial structures so that vaccines might be developed to disrupt their ability to reduce or eliminate *Salmonella* colonization [41-44]. While there are a variety of immunological responses to the fimbriae in the host, the responses that are directed against intraintestinal organisms appear to be ineffective and to date are unable to prevent the organism from colonizing the intestinal tract of the host [45, 46].

A wide variety of different factors play a role in the ability of *Salmonella* to establish itself as a long-term resident of the intestinal microbiome in poultry. Factors that impact the ability of the bacteria to become a microbiome resident include access to host cell receptors for anchoring of the organisms to the epithelium of the gut, competition for nutrients with other organisms in the microbiota for growth, and the effects of inhibitory substances (*i*.*e*., – bacteriocins, secondary metabolites, *etc*.) that are produced in the intestinal that inhibit growth of, or kill, *Salmonella*. Collectively, these factors are termed the “microbial barrier” that provides colonization resistance to organisms that may be pathogenic or deleterious to the host [47-49]. A prime example of the importance of the microbial barrier is observed when antibiotic treatment destroys the intestinal microbiome allowing *Clostridium difficile* opportunity to cause intestinal colitis [50]. Similarly, infection experiments performed in germ-free animals provide strong evidence that the ability of a pathogen to cause disease is significantly reduced by the presence of an intact microbiome [51, 52].

This work reports the characterization of an engineered probiotic derived from *E. coli* Nissle 1917 that expresses *p*-specific hyperadherence mediated by type 1 fimbriae (T1F), thereby exploiting the principles of colonization resistance to competitively exclude pathogenic *Salmonella* strains from the poultry intestinal tract. Unlike conventional antimicrobial strategies that attempt to eliminate pathogens after colonization has occurred, this approach is designed to occupy the ecological niche required for *Salmonella* persistence before pathogenic strains can establish themselves within the intestinal microbiome. The historical replacement of *S. Gallinarum* by *S. Enteriditis* in poultry production systems demonstrates that superior intestinal competitors can profoundly shape pathogen ecology and disease emergence. Our findings suggest that this engineered probiotic may function as a non-pathogenic competitive strain capable of displacing or preventing colonization by virulent *Salmonella* serovars through targeted niche occupation and adherence-based exclusion. As pressure increases to reduce antibiotic use in food animal production while simultaneously meeting increasingly stringent food safety standards, this strategy represents a potentially transformative platform technology for controlling enteric pathogens in poultry. Beyond reducing *Salmonella* carriage in flocks, this work establishes a foundation for the rational engineering of probiotics that leverage competitive microbial ecology to improve animal health, enhance food safety, and reduce the burden of foodborne disease transmission to humans.

## Materials and Methods

### Strains, Plasmids and Growth Conditions

*Salmonella enterica* serovar Typhimurium (*S*. Typhimurium) strains SL1344 [53] and DT104 [31] were used as representative pathogenic wild-type *Salmonella* strains and have been characterized previously. The *S*. Typhimurium BJ3714 Δ*fimA* and *S*. Typhimurium BJ3716 Δ*hilD*-URS Δ*ssaV* strains were created using the one-step inactivation mutagenesis technique, using plasmids pKD3, pKD4, pKD46 and pCP20 [32]. Mutant strains constructed with this procedure were confirmed to have the desired mutations by PCR analysis. *E. coli* Nissle 1917 was used as the scaffold strain for the engineered probiotic strain. The *fimA* and *asd* genes were deleted sequentially in *E. coli* Nissle 1917 using the one-step inactivation mutagenesis procedure [32]. The desired mutations were confirmed by PCR analysis and by functional assay for adherence or for a requirement for DAP in the growth media.

Plasmid pBDJ450 was constructed by cloning partially digested (Sau3A) chromosomal DNA into pACYC184 and identifying a clone that conferred adherence onto *E. coli* HB101, which was subsequently proven to be mannose-sensitive adherence. Plasmid pBDJ450 was modified by cloning the *E. coli asd* gene onto plasmid pBDJ450. The functional *asd* gene was inserted into orf stm0551 which is a repressor of *Salmonella* type 1 fimbrial expression [54]. The inserted *asd* was shown to complement an *E. coli* strain with an *asd* deletion and the plasmid was confirmed to have intact *S*. Typhimurium type 1 fimbrial genes.

Bacterial strains were grown on L agar with appropriate antibiotics, as needed. Antibiotics were added at the following concentrations ampicillin, 100 μg ml^−1^; kanamycin, 25 μg ml^−1^; chloramphenicol, 25 μg ml^−1^; tetracycline, 100 μg ml^−1^. Broth cultures of bacteria were grown statically in 10 mls of L broth with appropriate antibiotics at 37°C and passaged after 48 hours at least three times for maximal expression of type 1 fimbrial structures and binding activity.

### Yeast Agglutination Assay

*S. cerevisiae* yeast cells were grown with shaking overnight at 37°C in 100 mL of medium containing peptone (2%; Difco), yeast extract (1%), and glucose (2%). The cells were harvested by centrifugation at 1,500xg for 10 min, washed by resuspension in 100 mL of phosphate-buffered saline, pH 7.4, and sedimented again. The pellet was resuspended in 50 mL PBS with 0.2% glutaraldehyde and incubated at room temperature for 1 hour. The yeast cells were then pelleted at 1,500xg for 10 mins, washed twice with PBS and incubated in 50 mL of PBS with 10 mg/mL of glycine for 30 mins at room temperature. The fixed yeast cells were then washed again 2x in PBS and resuspended in 3 mL of PBS with 0.02% sodium azide to prevent bacterial growth and the cells could be stored up to 1 month at 4°C. A typical agglutination assay used 20 μL of the fixed yeast cells mixed with an equal volume (20 μL) of a suspension of a bacterial strain on a microscope slide and gently mixed to allow binding of the bacteria to the fixed yeast cells. A positive agglutination reaction gave clear clumping of the yeast cells that was mediated by binding and crosslinking via the bacteria. A negative reaction typically gave no clear clumping of yeast cells. Microscopic examination of a coverslip under low power was sometimes used to confirm agglutination reactions [55].

### Bacterial Adherence Assay

The adherence assay has previously been described by our group [12]. Briefly, HeLa cells were maintained in RPMI 1640 tissue culture media (Gibco) with 5% fetal calf serum (FCS). 10^5^ cells were seeded into each well of a 24-well tissue culture dish and incubated overnight. The following day 1×10^8^ CFU of bacteria were added to each well (ratio of bacteria to cell 1000:1) and the bacteria were allowed to bind to the tissue culture cells for 3 hours at 37°C in the tissue culture incubator with gentle agitation. Following the three-hour incubation, each well was washed 3x with 1 mL of sterile PBS to remove non-adherent organisms and organisms that might be loosely adherent to the tissue culture plastic. After a final wash, the cells were incubated for 10 mins at room temperature in 200 μL of PBS with 0.2% Triton X-100 to solubilize the cells and release the bacteria from the membranes of the host cells and then 800 μL of LB broth was added to each well to dilute the Triton X-100. Ten-fold dilutions from each well were then plated onto L agar with appropriate antibiotics to quantitate the number of organisms that had attached to each tissue culture monolayer over the course of the 3-hour assay. Percent adherence was calculated by determining the total number of CFU recovered from each well and divided by the input number of bacteria (1×10^8^ CFU). Adherence for the strains is expressed either as total CFU for a given assay or as percent adherence of the input.

For the competitive adherence assays, two strains were added to the same tissue culture well at the same time. All other steps for the adherence assay were as described above.

### Inhibition of type 1 fimbrial binding to *Salmonella* rabbit polyclonal antibody

The inhibition of yeast agglutination by the *Salmonella* anti-type 1 fimbrial polyclonal rabbit antibody was assessed by following the protocol for the agglutination assay but 5 μL of polyclonal antibody was added to the bacteria suspension before it was added to the yeast cells. All other procedures of the assay were the same as described previously.

### *In vivo* competitive exclusion studies

Animal studies were performed at Iowa State University with approval from the Institutional Animal Care and Use Committee. For broilers and layers, one-day old birds were obtained from Welp Hatchery. Birds were mixed genders and various breeds. Animal husbandry was performed as per our recent study [56]. On days 3, 4, and 5 after arrival, birds were orally inoculated with 1.2 x 10^5^ CFU of either *E. coli* Nissle 1917 pBDJ451 or *E. coli* Nissle 1917. On days 6, 7, and 8 after arrival, birds were orally inoculated with 1.2 x 10^5^ CFU of *Salmonella* Typhimurium strain LNWI [57]. Cloacal swabs (∼1gm) were then taken from each bird every three to four days and were subjected to enumerative *Salmonella* culture using XLD agar [57]. At the end of the study, birds were euthanized with a captive bolt and ceca were harvested for enumerative *Salmonella* culture [57]. Broiler and turkey study were performed on two separate occassions.

For layers, 20-week-old birds were housed in individual cages and procedures were performed as per the broiler studies except that the dose of inoculated bacteria was increased by ten-fold. Additionally, *S*. Enteriditis strain NVSL 1997 [31] was used as the test strain. For eggshell contamination studies, recovered eggs were soaked in 10 mL of LB broth of which 100 μL was plated on XLD agar for enumeration. Layer studies were performed once using each of the two *Salmonella* serovars. 20 birds were used in each group.

## Literature Cited

1. Falkow, S., Probing the intracellular life of bacteria. Harvey Lect, 1997. 93: p. 65–74.

2. Galan, J.E., Molecular genetic bases of Salmonella entry into host cells. Mol Microbiol, 1996. 20(2): p. 263–71.

3. Jones, B.D. and S. Falkow, Identification and characterization of a Salmonella typhimurium oxygen-regulated gene required for bacterial internalization. Infect Immun, 1994. 62(9): p. 3745–52.

4. Jones, B.D. and S. Falkow, Salmonellosis: host immune responses and bacterial virulence determinants. Annu Rev Immunol, 1996. 14: p. 533–61.

5. Hensel, M., Salmonella pathogenicity island 2. Mol Microbiol, 2000. 36(5): p. 1015–23.

6. Hensel, M., Evolution of pathogenicity islands of Salmonella enterica. Int J Med Microbiol, 2004. 294(2-3): p. 95–102.

7. Worley, M.J., Salmonella Type III Secretion System Effectors. Int J Mol Sci, 2025. 26(6).

8. dos Santos, A.M.P., R.G. Ferrari, and C.A. Conte-Junior, Virulence Factors in Salmonella Typhimurium: The Sagacity of a Bacterium. Current Microbiology, 2019. 76(6): p. 762–773.

9. Sia, C.M., et al., Salmonella pathogenicity islands in the genomic era. Trends Microbiol, 2025. 33(7): p. 752–764.

10. Barrow, P.A., et al., The long view: Salmonella--the last forty years. Avian Pathol, 2012. 41(5): p. 413–20.

11. Silva, C., E. Calva, and S. Maloy, One Health and Food-Borne Disease: Salmonella Transmission between Humans, Animals, and Plants. Microbiol Spectr, 2014. 2(1): p. OH–0020-2013.

12. Boddicker, J.D., et al., Differential binding to and biofilm formation on, HEp-2 cells by Salmonella enterica serovar Typhimurium is dependent upon allelic variation in the fimH gene of the fim gene cluster. Mol Microbiol, 2002. 45(5): p. 1255–65.

13. Ledeboer, N.A. and B.D. Jones, Exopolysaccharide sugars contribute to biofilm formation by Salmonella enterica serovar typhimurium on HEp-2 cells and chicken intestinal epithelium. J Bacteriol, 2005. 187(9): p. 3214–26.

14. Gerlach, G.F., et al., Expression of type 1 fimbriae and mannose-sensitive hemagglutinin by recombinant plasmids. Infect Immun, 1989. 57(3): p. 764–70.

15. Kolenda, R., M. Ugorski, and K. Grzymajlo, Everything You Always Wanted to Know About Salmonella Type 1 Fimbriae, but Were Afraid to Ask. Front Microbiol, 2019. 10: p. 1017.

16. Thiagarajan, D., H.L. Thacker, and A.M. Saeed, Experimental infection of laying hens with Salmonella enteritidis strains that express different types of fimbriae. Poult Sci, 1996. 75(11): p. 1365–72.

17. Wilson, R.L., et al., Salmonella enterica serovars gallinarum and pullorum expressing Salmonella enterica serovar typhimurium type 1 fimbriae exhibit increased invasiveness for mammalian cells. Infect Immun, 2000. 68(8): p. 4782–5.

18. Ledeboer, N.A., et al., Salmonella enterica serovar Typhimurium requires the Lpf, Pef, and Tafi fimbriae for biofilm formation on HEp-2 tissue culture cells and chicken intestinal epithelium. Infect Immun, 2006. 74(6): p. 3156–69.

19. Hancox, L.S., K.S. Yeh, and S. Clegg, Construction and characterization of type 1 non-fimbriate and non-adhesive mutants of Salmonella typhimurium. FEMS Immunol Med Microbiol, 1997. 19(4): p. 289–96.

20. Old, D.C. and J.P. Duguid, Selective outgrowth of fimbriate bacteria in static liquid medium. J Bacteriol, 1970. 103(2): p. 447–56.

21. Firon, N., I. Ofek, and N. Sharon, Carbohydrate-binding sites of the mannose-specific fimbrial lectins of enterobacteria. Infect Immun, 1984. 43(3): p. 1088–90.

22. Hultgren, S.J., S. Normark, and S.N. Abraham, Chaperone-assisted assembly and molecular architecture of adhesive pili. Annu Rev Microbiol, 1991. 45: p. 383–415.

23. Sokurenko, E.V., et al., FimH family of type 1 fimbrial adhesins: functional heterogeneity due to minor sequence variations among fimH genes. J Bacteriol, 1994. 176(3): p. 748–55.

24. Sokurenko, E.V., et al., Quantitative differences in adhesiveness of type 1 fimbriated Escherichia coli due to structural differences in fimH genes. J Bacteriol, 1995. 177(13): p. 3680–6.

25. Rabsch, W., et al., Competitive exclusion of Salmonella enteritidis by Salmonella gallinarum in poultry. Emerg Infect Dis, 2000. 6(5): p. 443–8.

26. Lee, C.A., B.D. Jones, and S. Falkow, Identification of a Salmonella typhimurium invasion locus by selection for hyperinvasive mutants. Proc Natl Acad Sci U S A, 1992. 89(5): p. 1847–51.

27. Baxter, M.A., et al., HilE interacts with HilD and negatively regulates hilA transcription and expression of the Salmonella enterica serovar Typhimurium invasive phenotype. Infect Immun, 2003. 71(3): p. 1295–305.

28. Catron, D.M., et al., The Salmonella-containing vacuole is a major site of intracellular cholesterol accumulation and recruits the GPI-anchored protein CD55. Cell Microbiol, 2002. 4(6): p. 315–28.

29. Gallois, A., et al., Salmonella pathogenicity island 2-encoded type III secretion system mediates exclusion of NADPH oxidase assembly from the phagosomal membrane. J Immunol, 2001. 166(9): p. 5741–8.

30. Klein, J.R. and B.D. Jones, Salmonella pathogenicity island 2-encoded proteins SseC and SseD are essential for virulence and are substrates of the type III secretion system. Infect Immun, 2001. 69(2): p. 737–43.

31. Carlson, S.A., et al., Detection of multiresistant Salmonella typhimurium DT104 using multiplex and fluorogenic PCR. Mol Cell Probes, 1999. 13(3): p. 213–22.

32. Datsenko, K.A. and B.L. Wanner, One-step inactivation of chromosomal genes in Escherichia coli K-12 using PCR products. Proc Natl Acad Sci U S A, 2000. 97(12): p. 6640–5.

33. Kisiela, D., et al., Functional characterization of the FimH adhesin from Salmonella enterica serovar Enteritidis. Microbiology (Reading), 2006. 152(Pt 5): p. 1337–1346.

34. Kisiela, D., et al., Characterization of FimH adhesins expressed by Salmonella enterica serovar Gallinarum biovars Gallinarum and Pullorum: reconstitution of mannose-binding properties by single amino acid substitution. Infect Immun, 2005. 73(9): p. 6187–90.

35. CDC, https://www.cdc.gov/drugresistance/index.html. 2023.

36. Ruvalcaba-Gomez, J.M., et al., Non-Antibiotics Strategies to Control Salmonella Infection in Poultry. Animals (Basel), 2022. 12(1).

37. Clegg, S., L.S. Hancox, and K.S. Yeh, Salmonella typhimurium fimbrial phase variation and FimA expression. J Bacteriol, 1996. 178(2): p. 542–5.

38. Purcell, B.K., J. Pruckler, and S. Clegg, Nucleotide sequences of the genes encoding type 1 fimbrial subunits of Klebsiella pneumoniae and Salmonella typhimurium. J Bacteriol, 1987. 169(12): p. 5831–4.

39. Swenson, D.L. and S. Clegg, Identification of ancillary fim genes affecting fimA expression in Salmonella typhimurium. J Bacteriol, 1992. 174(23): p. 7697–704.

40. Tinker, J.K. and S. Clegg, Characterization of FimY as a coactivator of type 1 fimbrial expression in Salmonella enterica serovar Typhimurium. Infect Immun, 2000. 68(6): p. 3305–13.

41. De Buck, J., et al., Protection of laying hens against Salmonella Enteritidis by immunization with type 1 fimbriae. Vet Microbiol, 2005. 105(2): p. 93–101.

42. Musa, H.H., et al., The molecular adjuvant mC3d enhances the immunogenicity of FimA from type I fimbriae of Salmonella enterica serovar Enteritidis. J Microbiol Immunol Infect, 2014. 47(1): p. 57–62.

43. Duguid, J.P. and I. Campbell, Antigens of the type-1 fimbriae of salmonellae and other enterobacteria. J Med Microbiol, 1969. 2(4): p. 535–53.

44. Laniewski, P., et al., Analysis of Spleen-Induced Fimbria Production in Recombinant Attenuated Salmonella enterica Serovar Typhimurium Vaccine Strains. mBio, 2017. 8(4).

45. Curtiss, R., Vaccines to Control Salmonella in Poultry. Avian Dis, 2024. 67(4): p. 427–440.

46. Doyle, M.P. and M.C. Erickson, Reducing the carriage of foodborne pathogens in livestock and poultry. Poult Sci, 2006. 85(6): p. 960–73.

47. Acheson, D.W. and S. Luccioli, Microbial-gut interactions in health and disease. Mucosal immune responses. Best Pract Res Clin Gastroenterol, 2004. 18(2): p. 387–404.

48. Jones, B., L. Pascopella, and S. Falkow, Entry of microbes into the host: using M cells to break the mucosal barrier. Curr Opin Immunol, 1995. 7(4): p. 474–8.

49. Santos, R.L., et al., Life in the inflamed intestine, Salmonella style. Trends Microbiol, 2009. 17(11): p. 498–506.

50. Delmee, M. and M. Warny, Clostridium difficile colitis: recent therapeutical and immunological considerations. Acta Gastroenterol Belg, 1995. 58(3-4): p. 313–7.

51. Stecher, B., Establishing causality in Salmonella-microbiota-host interaction: The use of gnotobiotic mouse models and synthetic microbial communities. Int J Med Microbiol, 2021. 311(3): p. 151484.

52. Woelfel, S., M.S. Silva, and B. Stecher, Intestinal colonization resistance in the context of environmental, host, and microbial determinants. Cell Host Microbe, 2024. 32(6): p. 820–836.

53. Wray, C. and W.J. Sojka, Experimental Salmonella typhimurium infection in calves. Res Vet Sci, 1978. 25(2): p. 139–43.

54. Wang, K.C., et al., A previously uncharacterized gene stm0551 plays a repressive role in the regulation of type 1 fimbriae in Salmonella enterica serotype Typhimurium. BMC Microbiol, 2012. 12: p. 111.

55. Eshdat, Y., et al., Isolation of a mannose-specific lectin from Escherichia coli and its role in the adherence of the bacteria to epithelial cells. Biochem Biophys Res Commun, 1978. 85(4): p. 1551–9.

56. Anderson, K., Burin, R., Rebollo, M., Krushinskie, E., Dridi, S. and S. Carlson., Reduction of Cecal Colonization and Fecal Shedding of Salmonella Typhimurium in Broilers Fed Proprietary Zinc-or Manganese-Amino Acid Complexes. J. Appl. Poultry Res, 2024. 33(1): p. 100388.

57. Anderson, K., Matthew, L., Brewer, T., Rasmussen, M.A. and S.A. Carlson, Identification of Heritage Chicken Breeds with Diminished Susceptibility to Intestinal Colonization by Multiple Antibiotic-Resistant Salmonella spp. Livestock Science, 2015. 182: p. 34–37.

